# Puget predicts gene expression across cell types using sequence and 3D chromatin organization data

**DOI:** 10.1101/2025.11.19.689320

**Authors:** Shengqi Hang, Xiao Wang, Ghulam Murtaza, Anupama Jha, Bo Wen, Tangqi Fang, Justin Sanders, Sheng Wang, William Stafford Noble

## Abstract

Gene expression is governed by both linear DNA sequence and three-dimensional (3D) chromatin architecture. Most gene expression prediction models rely on sequence alone, thereby failing to capture structural context and to generalize to unseen cell types. We present Puget, a deep learning model that predicts cell type-specific gene expression from sequence and Hi-C data, which captures 3D chromatin organization. Puget pairs pretrained sequence and Hi-C encoders with a lightweight transformer decoder. Using paired Hi-C/RNA-seq from 36 human and 4 mouse biosamples, we evaluate Puget’s ability to generalize to held-out genes, held-out biosamples, and from human to mouse. Relative to a sequence-only baseline, Puget improves cross-biosample Pearson correlation by up to 25% on highly variable genes in training biosamples and, unlike the sequence-only model, generalizes to held-out biosamples and across species. In addition, *in silico* perturbation experiments show that Puget can prioritize experimentally validated enhancer-gene pairs. Together, these results highlight a generalizable approach for modeling gene expression from sequence and 3D chromatin organization.

## 1 Introduction

Gene expression dictates cellular identity [1], orchestrates developmental trajectories [2], and its dysregulation underlies many diseases [3–5]. A central problem in modern genomics is to understand how complex gene expression programs emerge from the interplay of DNA sequence and cell type-specific context, including histone modifications, chromatin accessibility, and three-dimensional (3D) genome organization. 3D organization can influence expression by partitioning the genome into compartments [6], insulating genes from regulators [7, 8], and facilitating long-range enhancer-promoter contacts [9]. However, despite all that we have learned about these general principles, the extent and nature of the influence of 3D structure on a given gene’s expression in a given cell type remains unresolved. Although experimental perturbations can probe this relationship, such experiments are often costly and labor-intensive [10–13]. A complementary strategy is to build quantitative models that take sequence and 3D chromatin organization information as input, enabling *in silico* assessment of how structural changes impact gene expression and providing hypotheses about regulatory mechanisms through model interpretation.

Most existing gene expression predictors use sequence-to-expression (S2E) frameworks, which use only DNA sequence as input and predict gene expression levels [14–25]. Despite strong within-cell-type performance, S2E models do not generalize to unseen cell types because DNA sequence is invariant. More importantly, these models cannot predict how 3D chromatin organization modulates expression.

To predict how 3D chromatin organization influences expression, a predictive model needs to take in 3D chromatin organization data as input. The Hi-C assay [6], which measures DNA–DNA contacts genome-wide, is particularly relevant because it provides a snapshot of 3D chromatin organization and reveals how different sequence elements interact to drive gene expression. Therefore, a first step toward deciphering the impact of chromatin organization on gene expression is to build a computational model that takes in sequence and Hi-C as input to predict gene expression. Existing multi-modal methods, such as Seq-GraphReg [26], have combined these two modalities. However, Seq-GraphReg is fundamentally designed to be cell type-restricted. Its sequence encoder is trained from scratch to predict epigenomic tracks within a single cell type, and this learned sequence embedding is then combined with Hi-C–derived graph to predict gene expression for that same cell type. This model design requires training a separate model for each new cell type, making the model unable to generalize to unseen cell types.

To overcome this limitation, our goal is to develop a single, generalizable model that can be trained jointly across many cell types and can generalize to unseen cell types. We hypothesize that one approach to achieving this generalizability is to build the model from powerful, pretrained representations of sequence and structure, rather than retraining from scratch for each cell type. To achieve this, we leverage the power of foundation models, which are large-scale models pretrained on extensive datasets that learn generalizable representations for downstream tasks [27].

For sequence, we leverage the encoder from the Enformer model [23], which is a CNN-transformer hybrid model pretrained to predict thousands of epigenomic tracks across many cell types, allowing it to learn a general vocabulary of regulatory sequence motifs. For Hi-C inputs, we use our recently developed HiCFoundation [28], a masked autoencoder pretrained on Hi-C contact maps from hundreds of biosamples (i.e., cell lines or primary tissues). This large-scale pretraining enables HiCFoundation to learn robust, generalizable chromatin representations that are resilient to the data sparsity often found in individual Hi-C experiments. HiCFoundation has been shown to transfer well beyond its pretraining distribution, including to predicting epigenomic tracks from Hi-C on unseen cell types. We reason that these two foundation models play complementary roles. The Enformer encoder provides a cell-type-invariant vocabulary of potential regulatory motifs, while HiCFoundation encoder provides a cell-type-specific representation of the 3D chromatin contacts between those motifs.

In this work, we introduce Puget (**P**redicting **U**nseen-cell-type **G**ene **E**xpression from sequence and **T**hree-dimensional chromatin organization), a deep learning model that pairs a frozen Enformer sequence encoder with a frozen HiCFoundation structural encoder and trains a lightweight fusion decoder to predict RNA-seq expression. We hypothesized that, by combining the representation power of two pretrained foundation model encoders, efficient fine-tuning of Puget’s fusion decoder will enable our model to accurately predict cell type-specific expression. We curate data from 36 human biosamples and four mouse biosamples with paired Hi-C/RNA-seq data for training the Puget fusion decoder and for benchmarking. We evaluate performance in three settings: on held-out genes in training biosamples, on held-out biosamples, and across species. Relative to a sequence-only baseline, Puget improves cross-biosample Pearson correlation by up to 0.15 on highly variable genes in training biosamples and, unlike the sequence-only baseline, generalizes to held-out biosamples and across species. This generalization capability extends to mechanistic interpretation: by performing *in silico* knockout of the Hi-C input at candidate enhancer sites for held-out genes within a held-out biosample, Puget’s predicted gene expression changes successfully prioritize enhancer-gene pairs previously validated by CRISPRi experiments. Together, these results mark a first step toward generalizable, 3D structure-informed models of gene regulation.

## 2 Results

### 2.1 Puget integrates DNA sequence and 3D chromatin organization for gene expression prediction

Puget uses a dual-encoder, single-decoder architecture to predict cell type-specific gene expression by combining 1D sequence and 3D chromatin information (**Fig. 1a**). The model is built from two powerful, pretrained encoders—Enformer (∼228M parameters) for sequence and HiCFoundation (∼305M parameters) for Hi-C—which both remain frozen during fine-tuning. These encoders feed their respective embeddings into a lightweight transformer-based decoder (∼19M parameters). This decoder first projects the 1D sequence embeddings into a 2D grid via an outer sum, which is then added element-wise to the 2D Hi-C patch embeddings. The outer sum of the sequence embeddings creates a cell type-invariant 2D map of all potential motif-pair interactions. The subsequent element-wise addition integrates this map with the cell type-specific chromatin contacts from the Hi-C embeddings, allowing the decoder to identify which interactions are structurally co-localized. This fused 2D representation is then flattened and processed by a standard transformer to predict a scalar gene expression value (see **Supplementary Fig. S1** for architectural details).

**Figure 1.**
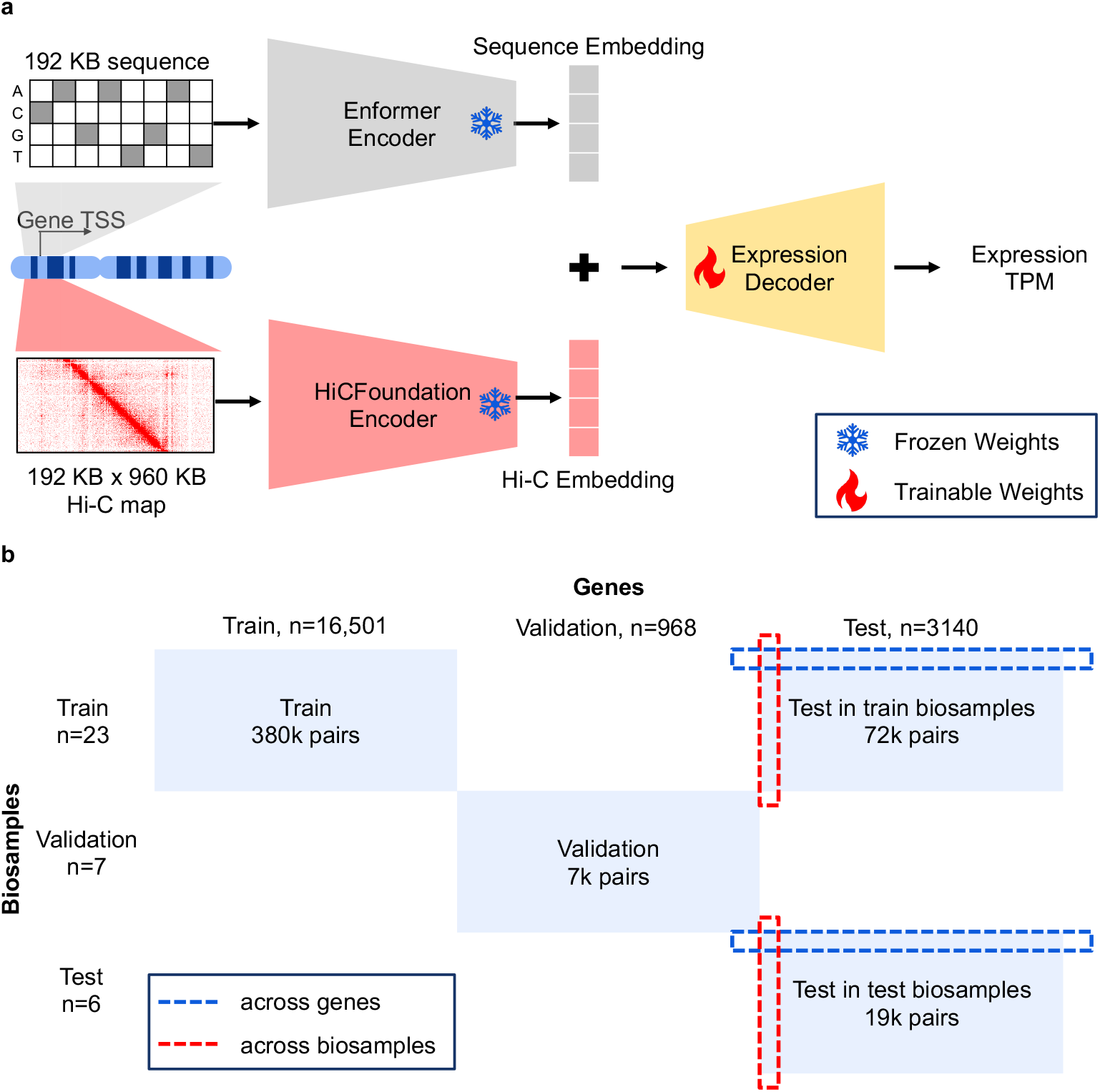
Puget model and data splits. **a** Schematic of Puget model. The inputs to the model are a 192 kb × DNA sequence and a 192 kb × 960 kb Hi-C submatrix, both centered at each gene’s transcriptional start site (TSS). The DNA sequence is processed by the pretrained Enformer model encoder (∼ 228M parameters), while the Hi-C submatrix is processed by the pretrained HiCFoundation model encoder (∼ 305M parameters). For gene expression fine-tuning, both encoders are frozen. Their output embeddings are then fed into a lightweight transformer decoder (∼ 19M parameters) to predict the log-transformed gene expression value. **b** Split of human biosample–gene pairs used to train, validate, and test gene expression prediction models. The data is split along both the gene and biosample axes. Model performance is evaluated using the Pearson correlation computed across genes and across biosamples. Data used for Enformer and HiCFoundation pretraining are not shown.

All inputs to the model are gene-centric, centered around each gene’s transcription start site (TSS). We provide a 192 kb DNA sequence to the sequence encoder and a 192 kb × 960 kb Hi-C submatrix at 1 kb resolution to the Hi-C encoder. The model is trained to predict scalar expression values derived from log-transformed transcripts-per-million (TPM) values.

To train and validate Puget, we curated a dataset of 36 human and four mouse biosamples with paired Hi-C and RNA-seq data publicly available from ENCODE [29]. We designed a rigorous evaluation framework to test model generalization at three levels: on held-out genes in training biosamples (**Fig. 1b**), on held-out genes in held-out biosamples (**Fig. 1b**), and across species from human to mouse. To prevent data leakage, all data splits were made consistent with the pretraining hold-outs of HiCFoundation (full details in **Methods**). Because the original Enformer model was not trained with this biosample hold-out scheme, we retrained an Enformer model using a subset of its pretraining data consistent with our split (retraining details in **Methods**). Puget uses the sequence encoder from this retrained model. Our data splitting scheme ensures that none of our test genes or test biosamples were seen during either encoder’s pretraining phase.

To contextualize Puget’s performance, we compared it against two single-modality baselines: a sequence-only baseline using the retrained Enformer and a Hi-C only baseline using HiCFoundation. Both baselines use the same respective frozen encoder paired with a fine-tuned transformer decoder. As in prior work [23, 24], we primarily assessed performance using the Pearson correlation between predicted and observed expression, both across genes and across biosamples (**Fig. 1b**).

### 2.2 Puget accurately predicts gene expression in seen biosamples

We first evaluated Puget in the standard setting for sequence-to-expression models: predicting expression for genes on held-out chromosomes within biosamples seen during training (**Fig. 2a**). This “seen-biosample, held-out chromosome” setting tests whether the model can generalize its learned regulatory grammar to unseen DNA sequences while remaining within familiar cellular context.

**Figure 2.**
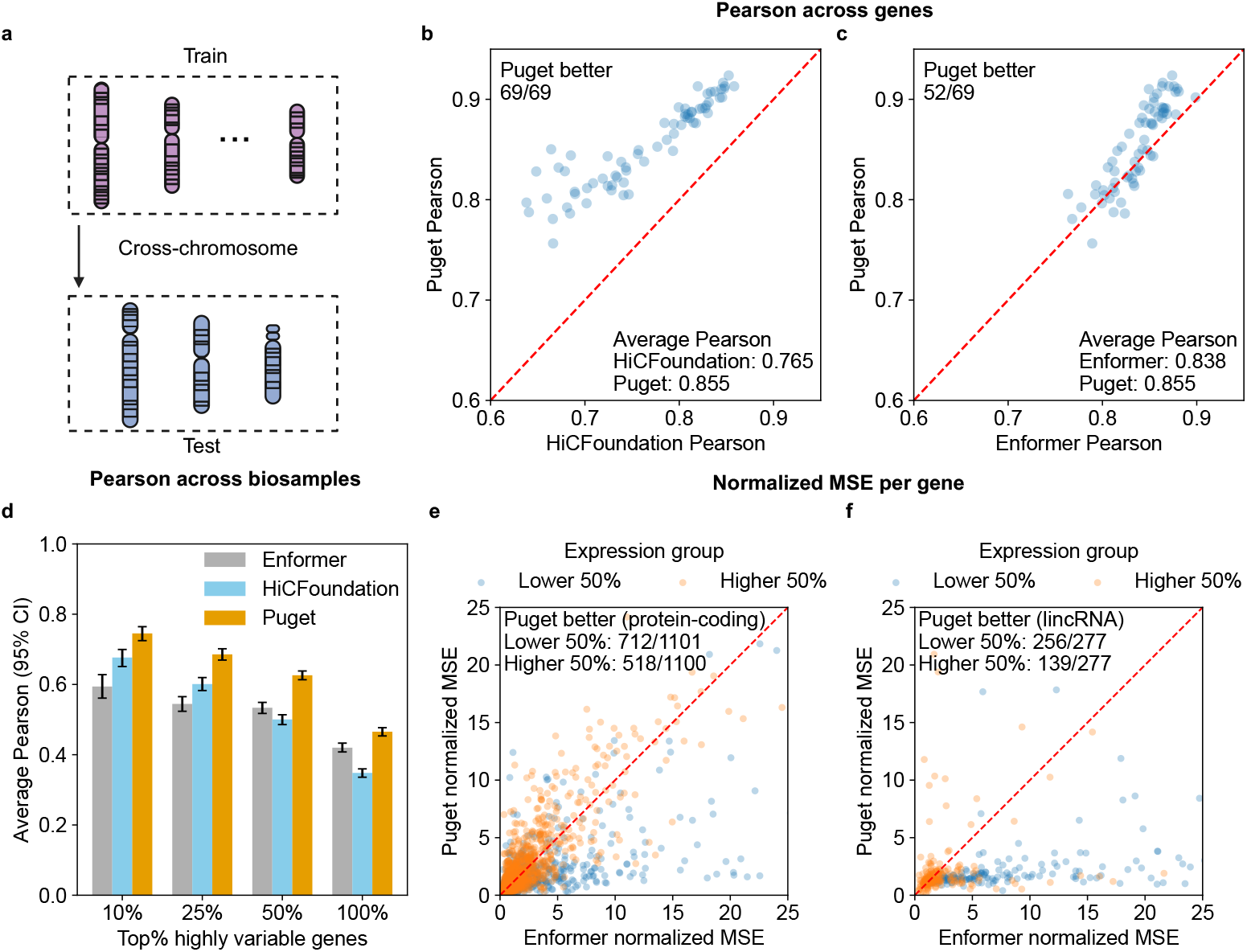
Puget accurately predicts gene expression in training (“seen”) biosamples. Unless noted, performance is Pearson correlation between observed and predicted log-expression. “Cross-gene” denotes correlation computed across annotated genes within a chromosome-biosample pair; “Cross-biosample” denotes correlation computed across biosamples for a single gene. **a** Schematic of the cross-chromosome setting in seen biosamples, where a selected set of chromosomes are held out for testing and the remaining chromosomes are used for training and validation. **b** Pairwise cross-gene performance comparing Puget and HiCFoundation for 23 training biosamples on three held-out test chromosomes (*n* = 69 chromosome-biosample pairs). Puget outperforms HiCFoundation in all 69 comparisons, with the mean Pearson correlation increasing from 0.765 to 0.855. **c** As in (b), versus Enformer; Puget outperforms Enformer in 52 of 69 comparisons, with the mean Pearson correlation increasing from 0.838 to 0.855. **d** Mean cross-biosample correlation as a function of the HVG threshold; across thresholds, Puget consistently surpasses both HiCFoundation and Enformer. Error bars indicate 95% confidence intervals estimated from 1,000 bootstrap resamples of genes at each threshold. **e** Scatterplot comparing variance-normalized mean squared error across training biosamples for Puget versus Enformer for each protein-coding gene with non-zero variance. Points are colored by average expression across training biosamples (top 50% vs. bottom 50%). For the higher 50%, Puget outperforms Enformer in 518 out of 1100 comparisons; for the lower 50%, Puget outperforms Enformer in 712 out of 1101 comparisons. **f** As in (e), for lincRNA genes. For the higher 50%, Puget outperforms Enformer in 139 out of 277 comparisons; for the lower 50%, Puget outperforms Enformer in 256 out of 277 comparisons. Schematic in **a** was created using BioRender (https://biorender.com).

In this setting, Puget outperformed both single-modality baselines. Across 69 test chromosomes from 23 human biosamples seen during training, Puget achieved a higher Pearson correlation than the HiCFoundation baseline on all 69 test chromosomes (sign-test p-value = 1.69 × 10^*−*21^, **Fig. 2b**) and surpassed the Enformer baseline on 52 of 69 chromosomes (sign-test p-value = 1.47 × 10^*−*5^, **Fig. 2c**). In terms of mean Pearson correlation, Puget reached 0.855, improving upon 0.765 for HiCFoundation and 0.838 for Enformer. These results suggest that jointly leveraging one-dimensional sequence and three-dimensional chromatin contacts enables Puget to more accurately capture regulatory signals than either sequence-only or Hi-C–only models.

Beyond predicting expression profiles within a biosample, a critical objective of an expression model is to capture cell-type specificity. Because cross-gene correlations can be dominated by ubiquitously expressed housekeeping genes, predicting the differential expression of cell type-specific genes is a more challenging task. Accordingly, for each test gene, we computed the Pearson correlation between observed and predicted expression across biosamples. For the set of test genes with the highest expression variance across biosamples (highly variable genes, HVGs), we then compared the mean of these per-gene correlations across models. At various thresholds for gene selection (top 10%, 25%, 50%, and 100% most variable genes, **Fig. 2d**), Puget consistently outperforms both the Enformer and HiCFoundation baselines, suggesting that Puget more effectively models biosample-specific expression patterns.

We further examined per-gene performance differences between Puget and Enformer using the mean squared error (MSE) across biosamples for each gene, normalized by the variance of each gene’s observed expression across biosamples. This variance-normalized error quantifies how well each model captures gene-specific expression patterns across conditions. We compared the performance separately for protein-coding and lincRNA genes in our test set, and we stratified genes by their average expression level across biosamples. For protein-coding and lincRNA genes (**Fig. 2e**,**f**), Puget performs comparably to Enformer for highly expressed genes and achieves lower normalized errors for lower-expressed genes.

We hypothesize that Puget shows limited gains for highly expressed protein-coding genes because many of these genes are broadly active across biosamples (for example, housekeeping genes) and are already well captured by sequence-only models. In contrast, many lowly expressed lincRNAs exhibit strong cell type-specificity [30] and have promoters with less conserved sequence motifs [31], making sequence alone an incomplete description of their regulation. In these cases, the Hi-C input provides cell type-specific chromatin interaction context that informs the regulation of these lincRNAs [32], enabling Puget to model their expression more accurately than a sequence-only approach. Consistent with this hypothesis, we observe a more pronounced improvement for Puget relative to Enformer for lowly expressed lincRNAs (**Fig. 2f**).

### 2.3 Puget generalizes to unseen biosamples

Although accurately modeling cell-type specificity within known biosamples is a key benchmark, the utility of a model also lies in its ability to generalize to new biological contexts to perform tasks such as gene expression prediction. This generalized gene expression prediction for unseen biosamples is essential for applications in novel cell types, tissues, or disease states where paired measured expression data is not available. We therefore tested Puget in this more stringent setting (**Fig. 3a**). Crucially, the test biosamples used in this evaluation were excluded from both the pretraining of the Enformer and HiCFoundation encoders and the fine-tuning of the Puget decoder. We hypothesized that the cell type-specific 3D chromatin information provided by the Hi-C modality would enable generalization to these unseen contexts.

**Figure 3.**
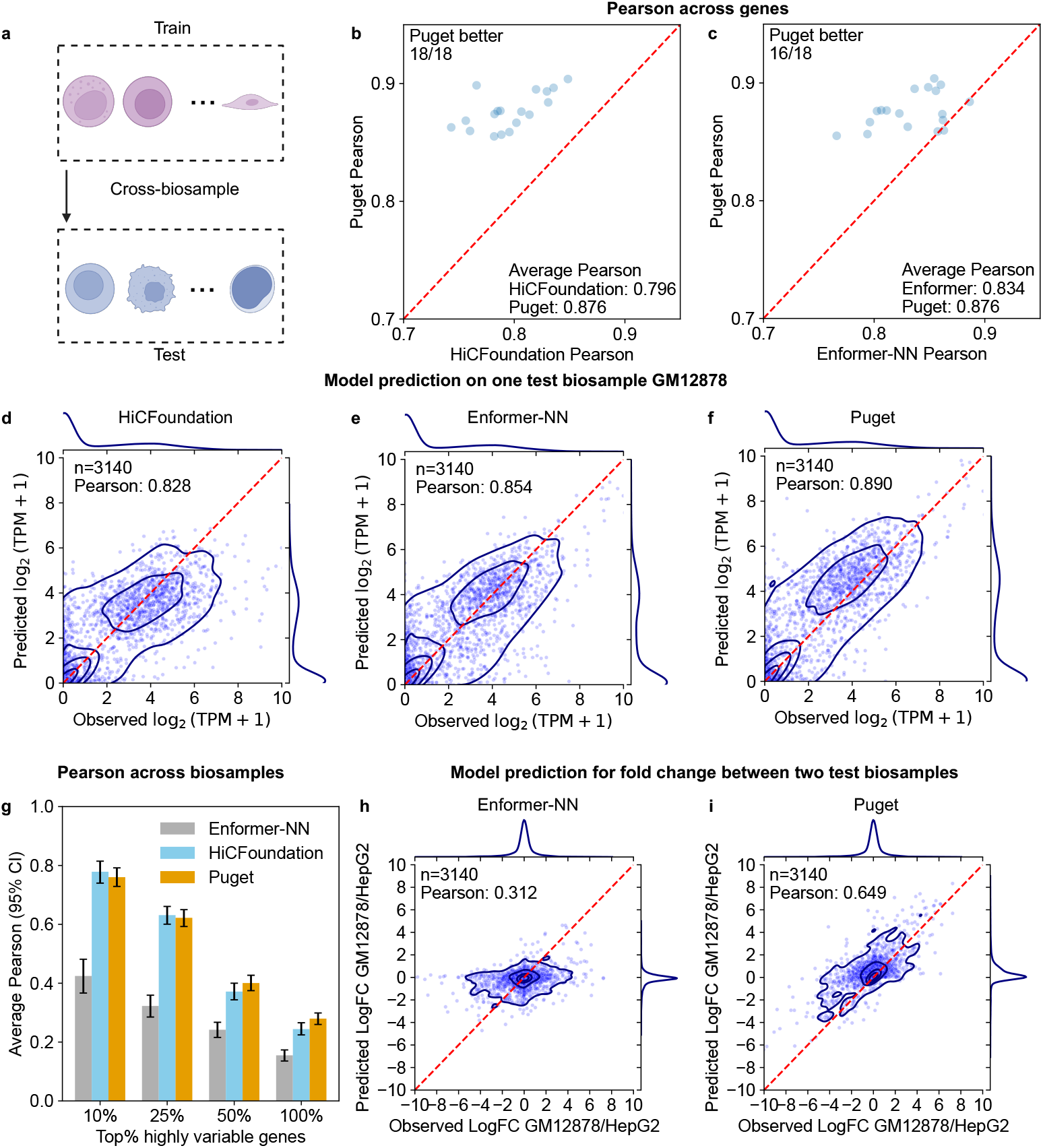
Puget generalizes to unseen biosamples. Enformer-NN imputes predictions for an unseen biosample by selecting its closest training biosample based on gene expression (see **Methods**) and using Enformer’s predictions for that matched biosample. **a** Schematic of the cross-biosample setting, in which a subset of biosamples is held out entirely for testing, while the training and validation chromosomes of the remaining biosamples are used for training and validation. **b** Pairwise cross-gene performance between Puget and HiCFoundation for six human test biosamples on three held-out test chromosomes (*n* = 18 chromosome-biosample pairs). Puget outperforms HiCFoundation in all 18 comparisons, with the mean Pearson correlation increasing from 0.796 to 0.876. **c** As in (b), versus Enformer-NN. Puget outperforms Enformer in 16 out of 18 comparisons, with the mean Pearson correlation increasing from 0.834 to 0.876. **d-f** Scatterplots of predicted versus observed TPM for one test biosample (GM12878) using HiCFoundation, Enformer-NN, and Puget, respectively. Puget attains the highest Pearson correlation. **g** Mean cross-biosample correlation as a function of the HVG threshold. Across thresholds, Puget consistently outperforms Enformer-NN and performs comparably with HiCFoundation. Error bars indicate 95% confidence intervals estimated from 1,000 bootstrap resamples of genes at each threshold. **h** Enformer-NN predicted versus observed log fold-change in expression between two test biosamples (GM12878 and HepG2) across genes on held-out chromosomes (each point is a gene). **i** Puget predicted versus observed log fold-change in expression between GM12878 and HepG2 across genes on held-out chromosomes. Puget shows higher agreement with observed fold-changes than Enformer-NN. Schematic in **a** was created using BioRender (https://biorender.com).

Because the Enformer baseline produces a fixed set of predictions corresponding to its training cell types, this model cannot directly predict expression for a new biosample that lacks a corresponding output head. Therefore, to construct a baseline for comparison, we developed an approach (“Enformer-NN”) that is based upon biosample nearest-neighbor matching. For a given test biosample, we computed the cosine similarity between the observed gene expression profile and each training biosample on the training genes, selected the most similar training biosample, and then used the Enformer baseline’s predictions for that matched biosample as the imputed prediction for the test biosample. This matching approach confers a significant advantage to the sequence-only baseline, because it uses the observed expression values of training genes in the test biosample, whereas Puget never has access to any expression data from the test biosamples. We also evaluated a variant of Enformer-NN that matches biosamples using similarities between Hi-C inputs, which performed worse than the expression-based matching approach (**Supplementary Fig. S2**).

In this challenging cross-biosample and cross-chromosome setting, Puget outperformed the HiCFoundation baseline and Enformer-NN baseline when performance was evaluated across genes. From 18 test chromosomes from six unseen human biosamples, Puget outperformed the HiCFoundation baseline in 18 out of 18 comparisons (sign-test p-value = 3.81 × 10^*−*6^, **Fig. 3b**) and outperformed the Enformer-NN baseline in 16 out of 18 comparisons (sign-test p-value = 6.56 × 10^*−*4^, **Fig. 3c**). Puget’s mean Pearson correlation of 0.876 improved upon corresponding values of 0.796 for HiCFoundation and 0.834 for Enformer. In one example test biosample GM12878, Puget also achieved the highest Pearson correlation compared to the HiCFoundation baseline and the Enformer-NN baseline (**Fig. 3d, e, f**).

In addition, when performance was evaluated across biosamples for highly variable genes, Puget consistently surpassed Enformer-NN at all gene selection thresholds (top 10%, 25%, 50%, 100%, **Fig. 3g**). This stronger tissue-specific predictive power is further demonstrated by Puget’s more accurate prediction of expression log fold-changes between two held-out cell lines, GM12878 and HepG2 (**Fig. 3h, i**). Notably, in this specific setting, Puget performed comparably to the HiCFoundation baseline (**Fig. 3g**), confirming that Hi-C is the primary driver of cell type-specific predictions. This performance comparison is not inconsistent with Puget’s advantage in the cross-gene analysis: predicting across genes requires capturing variation across genomic regions, where sequence contributes substantically and gives Puget an edge over the HiCFoundation baseline. In contrast, predicting across biosamples for a fixed gene relies on biosample-specific chromatin context, because the sequence is invariant; thus, in this setting, both Puget and HiCFoundation depend mainly on Hi-C and it becomes harder for Puget to outperform.

### 2.4 Puget provides an interpretable framework for analyzing sequence and 3D chromatin features in unseen biosamples

To investigate which features Puget uses when predicting expression in unseen biosamples, we applied two complementary interpretation strategies.

First, we used a gradient-based attribution method (input × gradient) for both sequence and Hi-C inputs [33]. For each held-out biosample, we computed saliency maps over the 192 kb sequence window and the corresponding 192 kb × 960 kb Hi-C submatrix and then averaged attributions across test genes. In both modalities, attributions peaked at bins overlapping the transcription start site (TSS), consistent with the importance of promoter-proximal sequence motifs and the promoter-enhancer contacts for gene regulation (**Supplementary Fig. S3**) [6, 9, 24].

Second, we performed a perturbation-based analysis to examine whether Puget can help prioritize active enhancers for genes. We used a compiled CRISPRi dataset of candidate enhancer–gene pairs in K562, one of our unseen test biosamples, containing both experimentally validated and non-validated enhancer– gene pairs [34]. For each enhancer candidate within the 960 kb window of a target gene, we held the sequence input fixed and perturbed only the Hi-C input by replacing the submatrix entries corresponding to the candidate enhancer region with a background distance-decay map estimated from K562 test regions (**Methods**). We then compared Puget’s predicted expression between the unperturbed (“wild-type”) and perturbed (“KO”) inputs and computed the magnitudes of the log fold-change values (**Fig. 4a**). Validated enhancers showed larger perturbation effects than non-validated candidates (**Fig. 4b**). However, the sign of the effect was not reliable (**Supplementary Fig. S4**). These results suggest that Puget can help prioritize candidate regulatory elements but should not be used to infer the direction of regulatory effects. Consistently, studies using S2E models also rely on absolute-valued perturbation or attribution scores to link enhancers to target genes [23–25]. Moreover, benchmarking studies of current S2E models [35, 36] show that S2E models frequently mis-predict the direction of genetic variant effects.

**Figure 4.**
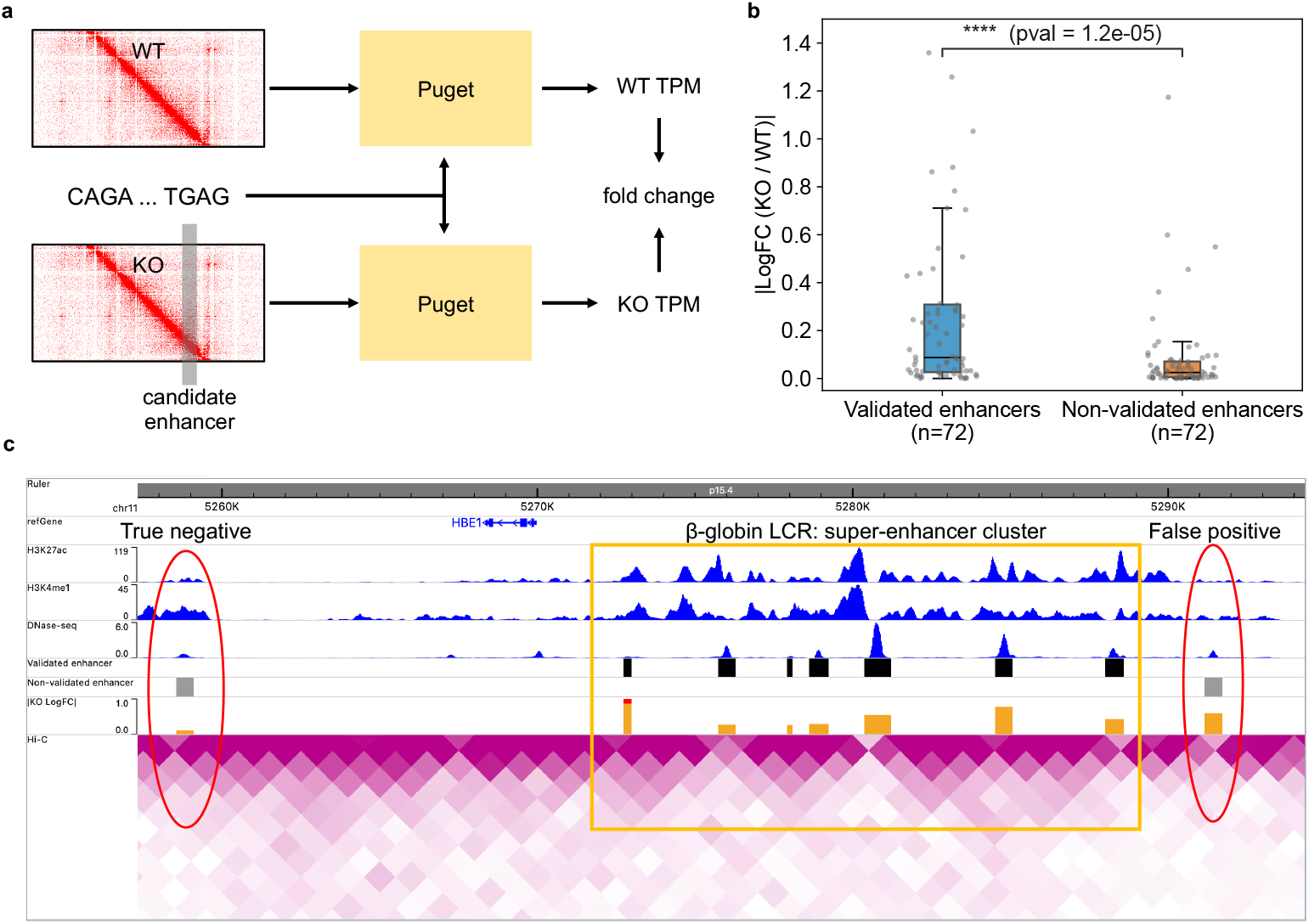
Puget provides an interpretable framework for analyzing sequence and 3D chromatin features in unseen biosamples. **a** Schematic of the *in silico* perturbation experiment. For a candidate enhancer-gene pair, the sequence input is held fixed, while the Hi-C input is perturbed by replacing a 3 kb × 192 kb region overlapping the candidate enhancer, with a background distance-decay map estimated from K562 test regions (**Methods**). Puget is applied to both the unperturbed (“WT”) input and the perturbed (“KO”) inputs, and the log fold-change between KO and WT predictions is used as a perturbation score. **b** Distribution of absolute fold-change for validated versus non-validated enhancer-gene pairs from the K562 CRISPRi dataset. Because the dataset contains many more non-validated pairs, we construct a distance-matched negative set by subsampling non-validated pairs to match the distance-to-TSS distribution of validated pairs. Validated enhancers show larger absolute effects than distance-matched non-validated enhancers. **c** Genome browser view of the *HBE1* locus. *In silico* perturbation produces large effects for DNase-hypersensitive sites within the *β*-globinn locus control region (LCR), while a downstream non-validated candidate shows only a small effect (“True negative”). A non-validated candidate near the LCR also exhibits a large fold-change (“False positive”).

We further examined a well-characterized locus, *HBE1*, in K562 [37, 38]. Puget assigned large absolute perturbation effects to multiple DNase-hypersensitive sites within the *β*-globin locus control region (LCR),consistent with their established roles as strong enhancers [37, 39, 40], while a downstream non-validated candidate showed minimal *in silico* perturbation effect (**Fig. 4c**). The model also produced one false positive near the super-enhancer cluster, potentially biased by the nearby strong chromatin signals rather than a true regulatory effect.

Overall, these analyses indicate that Puget can be used to interpret how sequence and 3D chromatin features contribute to its predictions, generating testable hypotheses about regulatory elements in unseen biosamples.

### 2.5 Puget generalizes from human to mouse

As a more rigorous test of generalization, we next assessed whether Puget learned fundamental regulatory principles that are conserved across species. To this end, we applied the human-trained model directly to a dataset of four mouse biosamples representing four distinct tissues (adrenal gland, gastrocnemius, left cerebral cortex, and heart) from the B6xCAST strain, without any fine-tuning (**Fig. 5a**). Not surprisingly, the HiCFoundation baseline showed reduced performance on mouse relative to human biosamples, with the mean Pearson correlation decreasing from 0.796 in human unseen biosamples (**Fig. 3b**) to 0.728 in mouse biosamples (**Fig. 5b**), reflecting species-specific nuances in the relationship between chromatin structure and transcription. Despite this challenging shift, Puget maintained a clear performance advantage over both single-modality baselines in cross-gene evaluations (**Fig. 5b, c**). To assess cross-tissue specificity, we quantified tissue-specific prediction by comparing predicted versus observed log fold-changes between all six unique tissue pairs for genes with non-constant expression across tissues. Puget outperformed HiCFoundation in two of six tissue pairs (non-overlapping 95% confidence intervals) and performed comparably in the remaining four (overlapping 95% confidence intervals) (**Fig. 5d**). This comparable performance between Puget and HiCFoundation in the cross-biosample setting is consistent with our observations in unseen human biosamples (**Fig. 3g**). Together, the cross-gene and cross-tissue results indicate that Puget captures conserved regulatory features that generalize across species.

**Figure 5.**
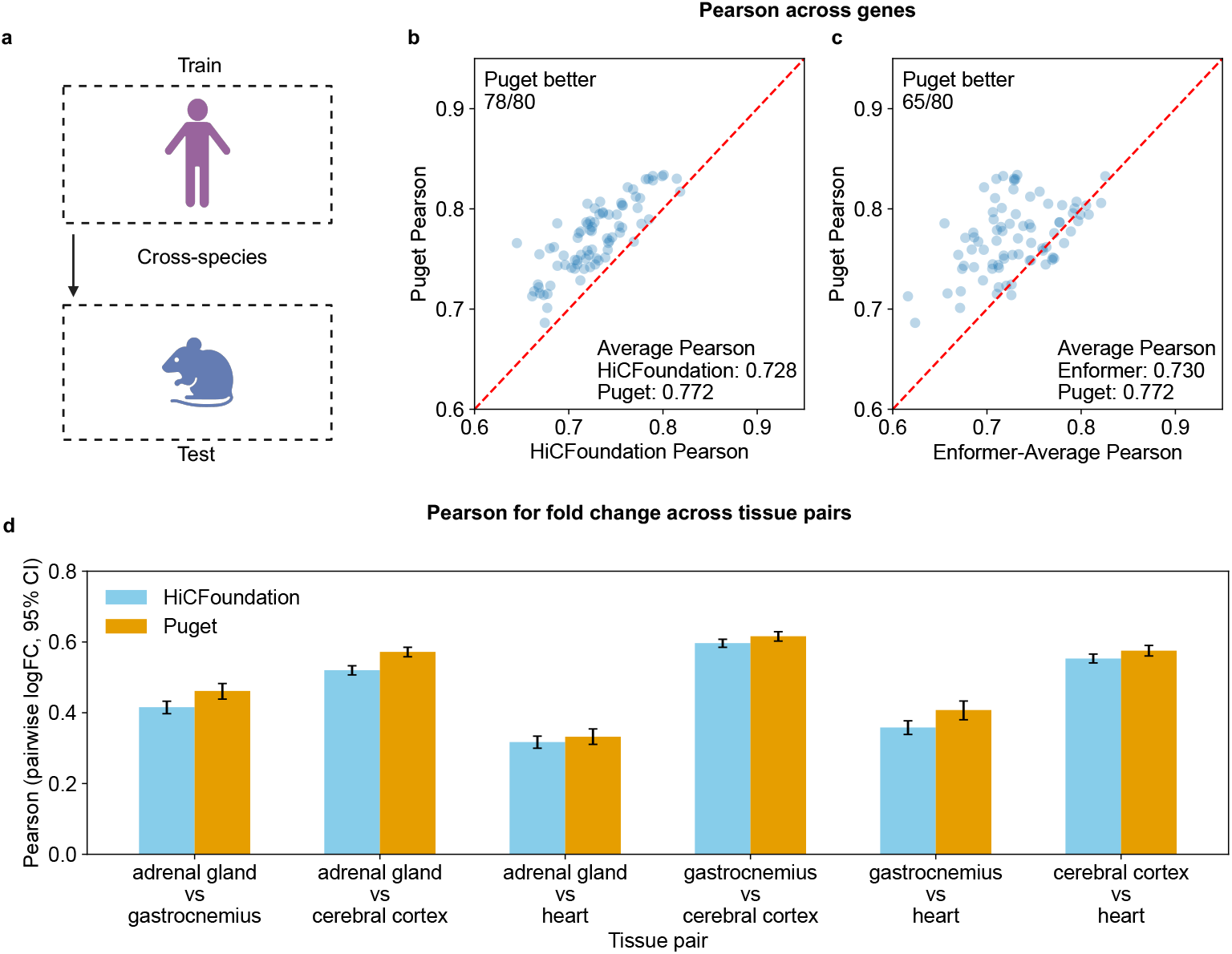
Puget generalizes from human to mouse. Enformer-Average applies the human-trained Enformer to mouse sequences by reusing the human biosample-specific output heads and averaging predictions across human biosamples to make a single prediction per mouse sequence (identical for all mouse biosamples). The mouse dataset includes four biosamples representing four distinct tissues (adrenal gland, gastrocnemius, left cerebral cortex, and heart). All four biosamples were used for the cross-gene evaluations in **b** and **c**, as well as the cross-tissue fold-change analysis in **d. a** Schematic of the cross-species setting, in which the model trained on human biosamples is directly applied to mouse biosamples without fine-tuning. **b** Pairwise cross-gene performance between Puget and HiCFoundation for four mouse biosamples on 20 mouse chromosomes (*n* = 80 chromosome–biosample pairs). Puget outperforms HiCFoundation in 78 out of 80 comparisons, with the mean Pearson correlation increasing from 0.728 to 0.772. **c** As in (b), versus Enformer-Average; Puget outperforms Enformer-Average in 65 of 80 comparisons, with the mean Pearson correlation increasing from 0.730 to 0.772. **d** Pearson correlation between predicted and observed log fold-changes for all six unique tissue pairs, computed on genes with non-constant expression across tissues. Error bars indicate 95% confidence intervals estimated from 1000 bootstrap resamples of genes. Schematic in **a** was created using BioRender (https://biorender.com).

## 3 Discussion

In this work, we introduce Puget, a multi-modal deep learning model that combines a frozen sequence encoder (Enformer) and a frozen Hi-C encoder (HiCFoundation) with a lightweight fusion decoder to predict gene expression from paired 1D sequence and 3D chromatin organization data. We evaluate generalization in three settings: (i) held-out chromosomes in training biosamples, (ii) held-out chromosomes in held-out biosamples, and (iii) zero-shot transfer from human to mouse, using two orthogonal metrics: per-biosample correlation across genes and per-gene correlation across biosamples. In held-out chromosomes of training biosamples, Puget outperforms both single-modality baselines on both metrics. In held-out-biosample and cross-species evaluations, Puget improves over sequence-only baselines for both metrics, and improves over Hi-C-only baselines for per-biosample (across-gene) correlation while performing comparably on per-gene (across-biosample) correlation. Finally, we demonstrate Puget’s utility for mechanistic interpretation in two ways: gradient-based attribution identifies the role of promoters in expression predictions, and *in silico* perturbation experiments prioritize enhancer-gene pairs validated by CRISPRi.

Puget extends recent efforts to move beyond purely sequence-based models toward structure-aware, biosample-specific prediction of gene expression [26, 41, 42]. Sequence-only models capture how DNA sequence shapes expression across genes but cannot directly account for biosample-specific chromatin context [19–25]. Previous structure-aware approaches have largely relied on 1D chromatin accessibility tracks [41, 42], limiting their ability to probe the role of 3D chromatin organization in transcriptional regulation. By using only sequence and Hi-C as inputs and reusing frozen, pretrained encoders with a small fusion decoder, Puget provides a simple, data-efficient, and compute-efficient approach that supports robust generalization across chromosomes, across biosamples, and from human to mouse.

An important nuance in our benchmarking results is that Puget outperforms HiCFoundation for per-biosample across-gene correlations in both training and unseen biosamples, and for per-gene across-biosample correlations in training biosamples, but shows comparable performance to HiCFoundation for per-gene across-biosample correlations in unseen biosamples. Conceptually, for each gene *j* in biosample *b*, we can approximate expression as,

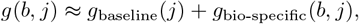

where *g*_baseline_(*j*) is largely sequence-driven and constant across biosamples, and *g*_bio-specific_(*b, j*) captures biosample-specific deviations that are influenced in part by chromatin organization. In the per-biosample across-gene setting, variation across genes is strongly shaped by *g*_baseline_(*j*), so a model like Puget that combines sequence inputs with Hi-C inputs consistently outperforms the Hi-C-only baseline. In the pergene across-biosample setting, however, *g*_baseline_(*j*) is constant and the correlation depends solely on how well a model captures *g*_bio-specific_(*b, j*) from biosample-specific features. Here, Puget performs better than HiCFoundation on training biosamples, but this advantage diminishes on unseen biosamples. A plausible explanation is that Puget’s mapping from Hi-C inputs to *g*_bio-specific_(*b, j*) is partially over-fitted to the distribution of training biosamples and generalizes less robustly to unseen chromatin contexts. Increasing the diversity of training biosamples or incorporating stronger regularization on this mapping may help improve out-of-distribution performance on this more challenging metric.

We anticipate several future directions to further improve the performance and utility of Puget. First, training and evaluating Puget on a larger and more diverse collection of paired Hi-C/RNA-seq datasets from ENCODE [29] and 4D Nucleome [43] consortia may improve robustness, particularly for unseen biosamples. In parallel, incorporating biosample-specific genomic sequences, rather than relying solely on the reference genome for cancer and highly mutated cell lines, could help the model better capture variant-dependent regulatory effects. Second, more systematic interpretability analyses are needed. Promising directions include using attribution methods to study interactions among pairs of sequence motifs, and exploring alternative perturbation schemes to probe the impact of perturbing specific loops, TADs, or compartments on predicted expression. Extending Puget to predict normalized expression profiles along the genome, rather than only gene-level TPM values, may also provide interpretation at finer resolution [44]. Third, Hi-C alone provides an incomplete and relatively coarse view of chromatin state. Future work could extend Puget to accept additional input modalities such as ATAC-seq or DNase-seq to obtain a more comprehensive representation of the cell type-specific regulatory landscape. Crucially, Puget’s modular architecture is extensible to this setting, allowing for the integration of additional frozen, pretrained encoders for these epigenetic modalities to refine predictions without training from scratch.

Puget provides a framework that reuses large, pretrained, modality-specific encoders with a lightweight fusion decoder to learn generalizable principles of how 1D sequence and 3D chromatin organization jointly contribute to gene expression across genes, biosamples, and species. We anticipate that such generalizable, multi-modal approaches will be increasingly useful for probing the regulatory impact of chromatin architecture and for generating testable hypotheses about transcriptional control.

## 4 Methods

### 4.2 Paired Hi-C and RNA-seq Datasets

The inputs to our model are the DNA sequence around each gene’s transcriptional start site (TSS) and a Hi-C submatrix centered on that TSS. The output is a scalar gene expression value for the corresponding gene, derived from an RNA-seq experiment. For DNA sequence, we extracted regions from the hg38 reference genome for human and the mm10 reference genome for mouse.

We curated 40 paired Hi-C/total RNA-Seq datasets from 40 distinct biosamples (36 human, 4 mouse) from the ENCODE [29] and 4D Nucleome [43] consortia (details, including accession numbers, are listed in **Supplementary Table 1**). “Paired” means that both assays derive from the same biosample, where “biosample” is an ENCODE term for a cell line, primary cell, or tissue.

All human and mouse Hi-C maps were generated by the intact Hi-C protocol developed by the Aiden lab at Baylor College of Medicine. We used unnormalized *cis* contact Hi-C maps at 1 kb resolution. All RNA-seq experiments were total RNA-seq produced by the Wold Lab at Caltech. We obtained transcripts-per-million (TPM) values from the ENCODE processing pipeline (RSEM v1.2.31) and log-transformed the values prior to modeling.

For human, we used the set of protein-coding and long intergenic noncoding RNA (lincRNA) genes from the ExPecto study [20], together with their CAGE representative TSS annotations (lifted over from hg19 to hg38) as defined in that work. For mouse, we started from protein-coding and lincRNA genes in GENCODE release M21 and followed the ExPecto protocol to select a CAGE representative TSS per gene. In both species, we excluded genes whose TSS lies within 2 Mb of a chromosome boundary. We further filtered by mappability and Hi-C coverage: a gene was removed if either (i) *>* 50% of bases within the 192 kb sequence window around its TSS are unmappable, or (ii) *>* 50% of Hi-C map rows intersecting the 192 kb region around its TSS have zero counts in the Hi-C maps aggregated across all training biosamples. After filtering, we have in total 20,609 human genes and 24,412 mouse genes.

We split the data along both biosample and gene axes, following the splits used by HiCFoundation, to avoid data leakage. Along the biosample axis, human biosamples were divided into 23 for training, 7 for validation, and 6 for testing, matching the HiCFoundation validation/test sets; all 4 mouse biosamples were reserved for testing only. Along the gene axis, genes on chromosomes 5, 11, and 14 were held out for testing, with chromosome 4 used for validation and all remaining chromosomes used for training. For human, this yields 16,501 training genes, 968 validation genes, and 3140 test genes. All mouse genes were reserved for testing.

To reflect the strand-specific nature of RNA-seq measurements, we orient all inputs in the direction of transcription (5’ to 3’). For genes on the reverse strand, we use the reverse complement of the 192 kb DNA sequence and reflect the Hi-C submatrix across both axes around the TSS so that the upstream and downstream contacts are aligned consistently with those of forward-strand genes.

### 4.2 Model architecture

#### 4.2.1 Hi-C map encoder

For input to the Hi-C encoder, we extract a 192 kb × 960 kb Hi-C input submatrix at 1 kb resolution centered on each gene’s TSS. The rectangular window captures distal enhancer–promoter contacts while avoiding the cost of a full 960 kb × 960 kb square. Hi-C features are computed using the pretrained HiCFoundation encoder [28], which we keep frozen while training our model.

HiCFoundation is a self-supervised masked autoencoder trained on millions of Hi-C submatrices. It treats each contact map as an image partitioned into 16 × 16 patches and randomly masks 75% of patches during pretraining. The encoder embeds unmasked patches (assigning a shared mask token to masked patches), and a lightweight decoder reconstructs the full map. The encoder consists of 24 transformer blocks with a latent dimension of 1024 (304 M parameters), while the decoder has 8 transformer blocks with a latent dimension of 512 (26 M parameters). Pretraining uses 116 million submatrices drawn from training chromosomes across 81 biosamples.

At 1 kb resolution, each patch spans 16 kb × 16 kb. Thus, our 192 kb × 960 kb window forms a 12 × 60 grid of patches. We interpolate the encoder’s 2D sinusoidal positional embeddings to this grid and pass all patches through the frozen encoder to obtain contextualized patch embeddings. We further take a center-crop of the embeddings and use the central 6 × 60 × 1024 embedding outputs from the frozen encoder to serve as the Hi-C representation for each gene. This region is fed into the decoder.

#### 4.2.2 DNA sequence encoder

Our DNA sequence encoder adopts the Enformer architecture [23], which uses learnable convolutional filters to detect local sequence motifs and transformer self-attention blocks to capture long-range dependencies. We do not use publicly released Enformer weights because the original pretraining did not hold out biosamples, raising potential data-leakage concerns for our evaluations. Instead, we retrain an Enformer backbone under the same chromosome/biosample split used for HiCFoundation pretraining. Starting from the original Enformer training set, we retain only human training examples, remove all biosamples reserved for validation or testing by HiCFoundation, omit all test chromosomes (chromosomes 4, 5, 11, 14) from training, and reserve chromosomes 9 and 10 for validation. This train/test split ensures that no sequences or biosamples used for evaluation are seen during sequence-encoder pretraining.

Following [23], we train the sequence encoder in a multi-task learning fashion: the input is a 196 kb sequence centered at the TSS, and the outputs are thousands of epigenomic tracks at 128 bp resolution. After training, we discard the final prediction head and freeze the remaining network as our sequence encoder. Applied to the 192 kb input, the encoder produces contextualized embeddings for 896 non-overlapping 128 bp bins covering the central 114 kb of the input. For downstream expression modeling, we extract the 750 bins (96 kb) centered on the TSS and pass their embeddings (750 × 3072) to the decoder as the sequence features.

#### 4.2.3 Puget gene expression decoder

The Puget decoder fuses sequence features from the central 96 kb window with Hi-C features from the aligned 96 kb × 960 kb submatrix (both centered at the TSS), and outputs a single scalar.

We first project modality-specific embeddings to a common channel dimension of 512 using two-layer MLPs. Sequence embeddings of shape 750 × 3072 (750 bins at 128 bp) are mapped to 750 × 512. Hi-C patch embeddings of shape 6 × 60 × 1024 are mapped to 6× 60× 512. The Hi-C grid is patched at 16 kb ×16 kb, whereas the sequence is binned at 128 bp. To align sequence and Hi-C spatial scales, we pool the projected sequence embeddings along the 128 bp axis to 16 kb. We employ three strided 1D convolutional blocks (each with a pool ratio of 5) to reduce the sequence embeddings to embeddings of shape 6 × 512.

Inspired by Akita [45], we lift the 1D sequence embedding 6 × 512 into a 2D “interaction matrix.” Specifically, given row-wise sequence features **s** ∈ ℝ^6×512^, we form a center interaction map,

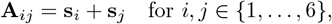

producing **A** ∈ ℝ^6×6×512^. We then zero-pad **A** along the distal dimension to match the Hi-C width, yielding **A**^*′*^∈ ℝ^6×60×512^.

We sum the projected Hi-C embedding and the padded sequence interaction map to obtain a fused embedding **F**∈ ℝ^6×60×512^. A 2D sinusoidal positional embedding is added to this fused grid, which is then flattened into a 1D sequence of 360 tokens. A learnable [CLS] token is prepended to this sequence, which is then processed by a lightweight standard transformer decoder of four transformer blocks. The final output representation of the [CLS] token is passed through a one-layer MLP to predict the final scalar gene expression value.

#### 4.2.4 Enformer gene expression decoder

For the sequence-only baseline, we keep the Enformer backbone frozen and attach a 1D decoder. First, a two-layer MLP projects the Enformer embeddings from 3072 to 512 channels. A learnable [CLS] token is prepended to the resulting 750 × 512 sequence embedding. A 1D transformer with four transformer blocks then processes this full sequence. We pass the final [CLS] token output representation to a single linear layer to predict gene expression for all training biosamples simultaneously, with one output head per biosample.

#### 4.2.5 HiCFoundation gene expression decoder

For the Hi-C-only baseline, we keep the HiCFoundation encoder frozen and attach a transformer decoder similar to Puget. A two-layer MLP projects Hi-C patch embeddings from 1024 to 512 channels, resulting in a 6 ×60× 512 embedding grid. Similar to the Puget decoder, a 2D positional embedding is added, and the grid is flattened into a 1D sequence. A learnable [CLS] token is prepended, and a standard transformer decoder of four transformer blocks processes the full sequence. The final output representation of the [CLS] token is passed to a single linear layer to output a scalar prediction.

## 4.3 Evaluation measures

We evaluated model performance in two orthogonal ways.

- **Across genes:** For each biosample, we computed the Pearson correlations between predicted and ground truth expression values over all test genes.
- **Across biosamples:** For each test gene *g*, we computed the Pearson correlation *r_g_* between predicted and ground truth values across biosamples. In this evaluation, we consider only the highly variable genes (top 10%, top 25%, top 50%, or top 100%). To select the highly variable genes, we first excluded genes with constant ground truth expression, ranked the remaining genes by variance, and selected the top percentage for evaluation. Our performance measure is the average correlation across these highly variable genes (HVGs), i.e., 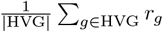.

## 4.4 Enformer baseline for unseen biosamples (Enformer-NN)

Because Enformer lacks biosample-specific heads for unseen biosamples, we impute its predictions via nearest-neighbor matching. For the results reported in the main text, we compute cosine similarity between each test biosample and all training biosamples using expression values on training genes. For each test biosample, we identify the most similar training biosample and use Enformer’s predictions for that closest match as the imputed predictions for the test biosample.

Alternatively, we also perform nearest-neightbor matching in Hi-C space. We consider two Hi-C similarity measures, averaging per-gene similarities over all genes in training chromosomes: (i) cosine similarity between HiCFoundation encoder embeddings of the gene-centric submatrix, and (ii) cosine similarity between PCA eigenvectors computed from the 960 kb region centered at the gene TSS. Cosine similarity on HiCFoundation embeddings performs better than that on the PCA-based eigenvectors, but both Hi-C based approaches underperform the expression-based matching.

### 4.5 Model training

#### 4.5.1 Enformer Retraining

We trained the Enformer model using Poisson negative log-likelihood loss between the predicted and ground truth epigenomic tracks. To reduce overfitting, we applied data augmentation by randomly shifting each input sequence by *±*3 bp and randomly reverse-complementing the sequence and correspondingly reversing its target tracks. We used the AdamW optimizer with default *β*_1_ = 0.9 and *β*_2_ = 0.999, and weight decay of 1× 10^*−*4^. The training was performed on multiple GPUs and with gradient accumulation, using an effective batch size of 64. The learning rate was linearly warmed up from 0 to 2 × 10^*−*4^ over the first five epochs, held constant for 80 epochs, and then decayed with a cosine schedule to 1 × 10^*−*6^ over the final 25 epochs. We picked the checkpoint with the lowest validation loss. Full training took approximately 4 days on 4 A100 GPUs (80 GB each). The retrained Enformer exhibits a moderate decrease in gene expression prediction performance relative to the public Enformer checkpoint on shared training biosamples (**Supplementary Fig. S5**). This reduction is expected given that the retrained model was trained on fewer training sequences, fewer epigenomic tracks, and only human data rather than both human and mouse.

#### 4.5.2 Decoder Training

To train Puget, we froze both the sequence and Hi-C encoders and updated only the decoders, optimizing with MSE loss. To reduce computational cost, we first computed and stored encoder embeddings for all inputs and then trained the decoder on these fixed embeddings. We used the Adam optimizer with default *β*_1_ = 0.9 and *β*_2_ = 0.999, and weight decay of 1× 10^*−*4^ with batch size 64. The learning rate was linearly warmed up from 0 to 1 × 10^*−*4^ over 2 epochs, then followed a cosine decay to 1× 10^*−*7^ over 18 epochs. We picked the checkpoint with the lowest validation loss. Under this setup, the decoder training took 8 hours on one A5000 GPU (24 GB).

For training single-modality models, we similarly froze the encoders and updated only the decoders, optimizing with MSE loss. Similarly, we first computed and stored encoder embeddings for all inputs and then trained the decoder on these fixed embeddings. We used Adam optimizer with default *β*_1_ = 0.9 and *β*_2_ = 0.999, and weight decay of 1 × 10^*−*4^ with batch size 64 for HiCFoundation and batch size 32 for Enformer. The learning rate was linearly warmed up from 0 to 1 × 10^*−*4^ over 2 epochs, then followed a cosine decay to 1 × 10^*−*7^ over 18 epochs. We picked the checkpoint with the lowest validation loss. Under this setup, the decoder training took 5 hours for HiCFoundation and 30 min for Enformer on one A5000 GPU (24 GB).

## 4.6 Model interpretation

To identify sequence and 3D chromatin contact features driving gene expression predictions of Puget, we apply input gradients (gradient × input) [33]. For Puget, gradients are computed with respect to the model inputs: (i) the one-hot DNA sequence and (ii) the Hi-C contact map at 1 kb resolution, both within a 192 kb × 960 kb window. Input gradient produces a base-resolution attribution map over the input sequence and an attribution matrix at 1 kb × 1 kb resolution for the input Hi-C.

For perturbation-based analysis of candidate enhancer–gene pairs, we kept the input sequence fixed and perturbed only the Hi-C input. Specifically, for a candidate enhancer located within the 960 kb window of a target gene, we masked a 192 kb × 3 kb column centered on the enhancer by replacing its Hi-C entries with values from a distance-decay-based expected matrix. To construct that expected matrix in K562, we pooled all test Hi-C submatrices, binned contacts by diagonal offset (genomic separation), and for each diagonal replaced entries with the median contact value for that diagonal. This procedure preserves the distance-based decay while removing locus-specific contact patterns at the perturbed region.

## 5 Data availability

All paired Hi-C and RNA-seq datasets used for training and benchmarking Puget were obtained from ENCODE portal. A complete list of ENCODE accession numbers is provided in **Supplementary Table 1**.

## 6 Code availability

The Puget software is open-source under an Apache 2.0 License. The code for data processing, model training and model inference is publicly available on Github at (https://github.com/Noble-Lab/Puget).

## 7 Acknowledgements

We thank Xinming Tu and Vikram Agarwal for helpful discussions and valuable feedback.

## Author contributions

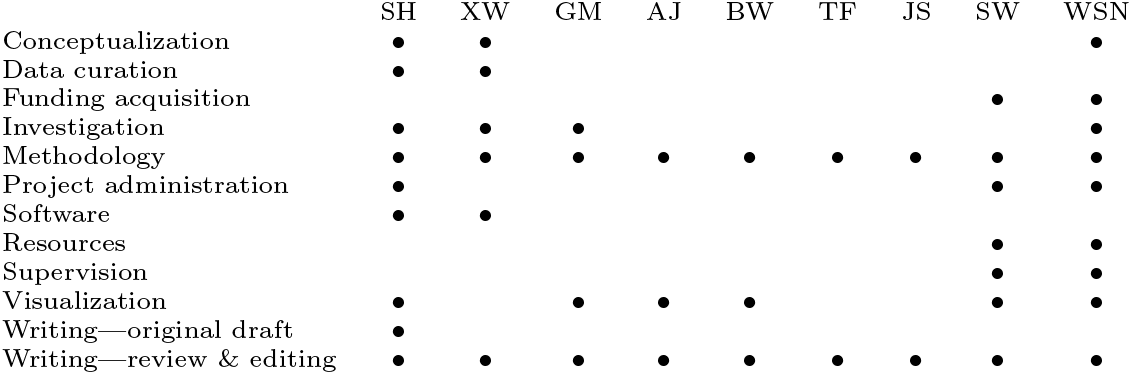

## Notes

### Competing Interest Statement

The authors have declared no competing interest.

