## supplementary-figures for "Puget predicts gene expression across cell types using sequence and 3D chromatin organization data"

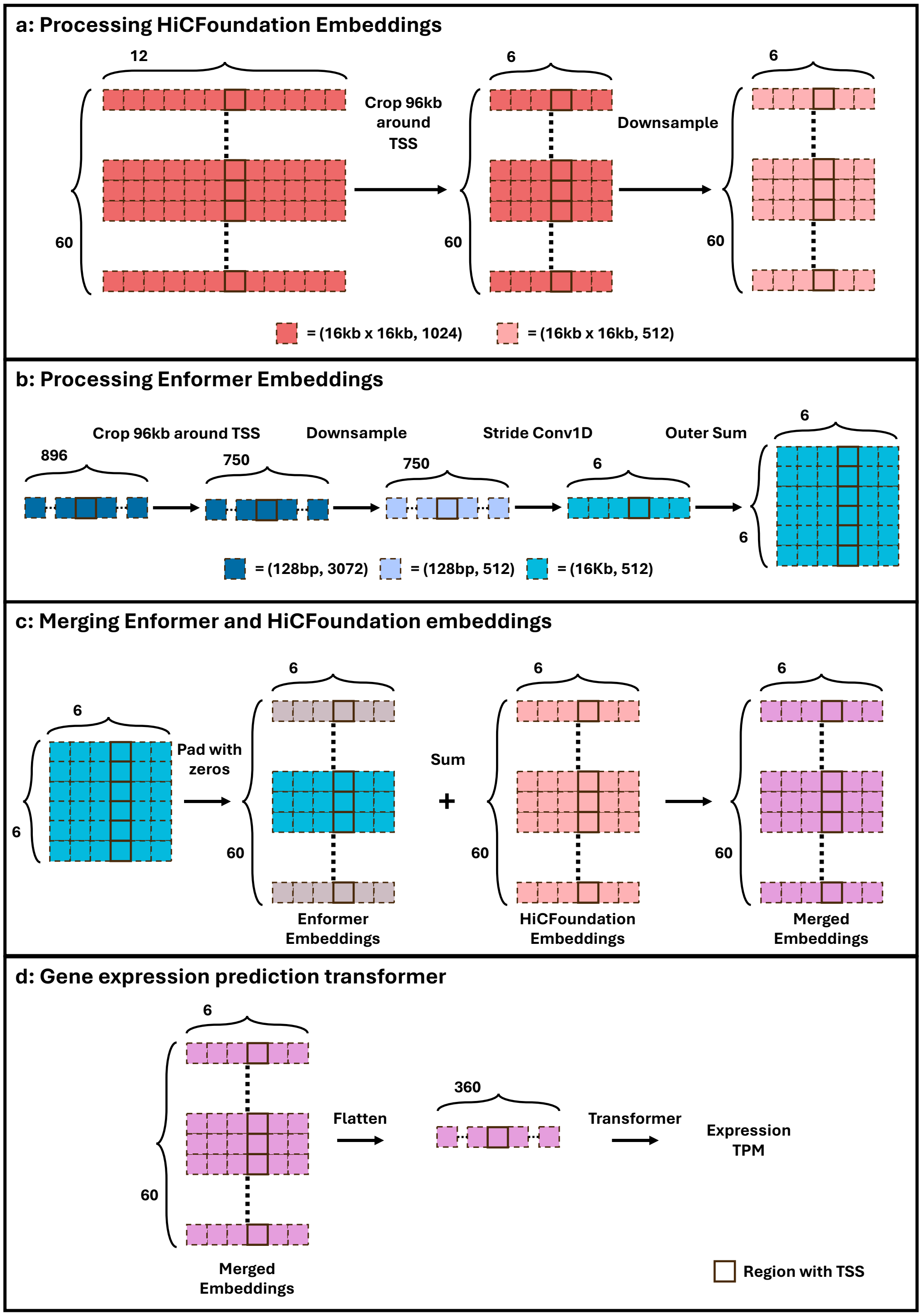


Figure S1: **Architecture of Puget Decoder.** **a** Hi-C embeddings from HiCFoundation encoder are center-cropped to a 96 kb × 960 kb window around the gene TSS from the original 192 kb × 960 kb window. **b** Sequence embeddings from Enformer encoder are center-cropped to a 96 kb window around the gene TSS from the original 114 kb embedding window. We apply convolutional pooling to downsample the sequence embedding to the same 16 kb resolution as the Hi-C embeddings, and then compute an outer sum to construct a 2D Enformer embedding grid. **c** We zero-pad additional rows in the Enformer 2D embedding grid to match the HiCFoundation 2D grid size and combine the two modalities by element-wise summation. **d** We add 2D sinusodial positional embeddings to the summed 2D grid, flatten the 2D grid to a sequence of embeddings, and pass the sequence through a transformer decoder to predict expression.

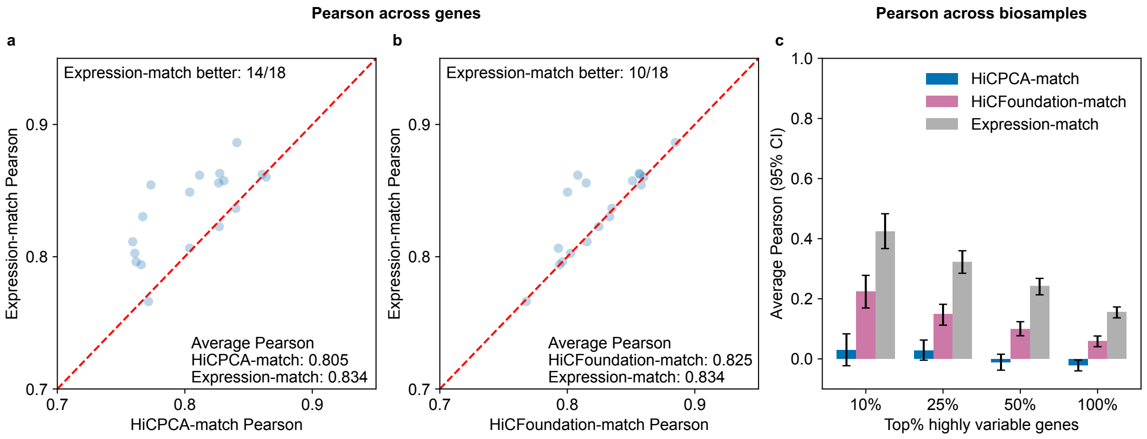


Figure S2: **Comparison between Enformer-NN variants using expression-based and Hi-C-based matching.** Enformer-NN imputes predictions for an unseen biosample by selecting its closest training biosample based on either gene-expression similarity or Hi-C–derived similarity (see Methods), and then using Enformer's predictions for that matched biosample. **a** Pairwise cross-gene performance between Enformer-NN with expression-match and Enformer-NN with HiCPCA-match for six human test biosamples on three held-out test chromosomes ($n$ = 18 chromosome-biosample pairs). The expression-match outperforms HiCPCA-match in 14 out of 18 comparisons, with the mean Pearson correlation increasing from 0.805 to 0.834. **b** Pairwise cross-gene performance between Enformer-NN with expression-match and Enformer-NN with HiCFoundation-match for six human test biosamples on three held-out test chromosomes ($n$ = 18 chromosome-biosample pairs). The expression-match outperforms HiCFoundation-match in 10 out of 18 comparisons, with the mean Pearson correlation increasing from 0.825 to 0.834. **c** Mean cross-biosample correlation as a function of the HVG threshold. Across thresholds, Enformer-NN with expression-match consistently outperforms Enformer-NN with HiCPCA-match and Enformer-NN with HiCFoundation-match. Error bars indicate 95% confidence intervals estimated from 1,000 bootstrap resamples of genes at each threshold.

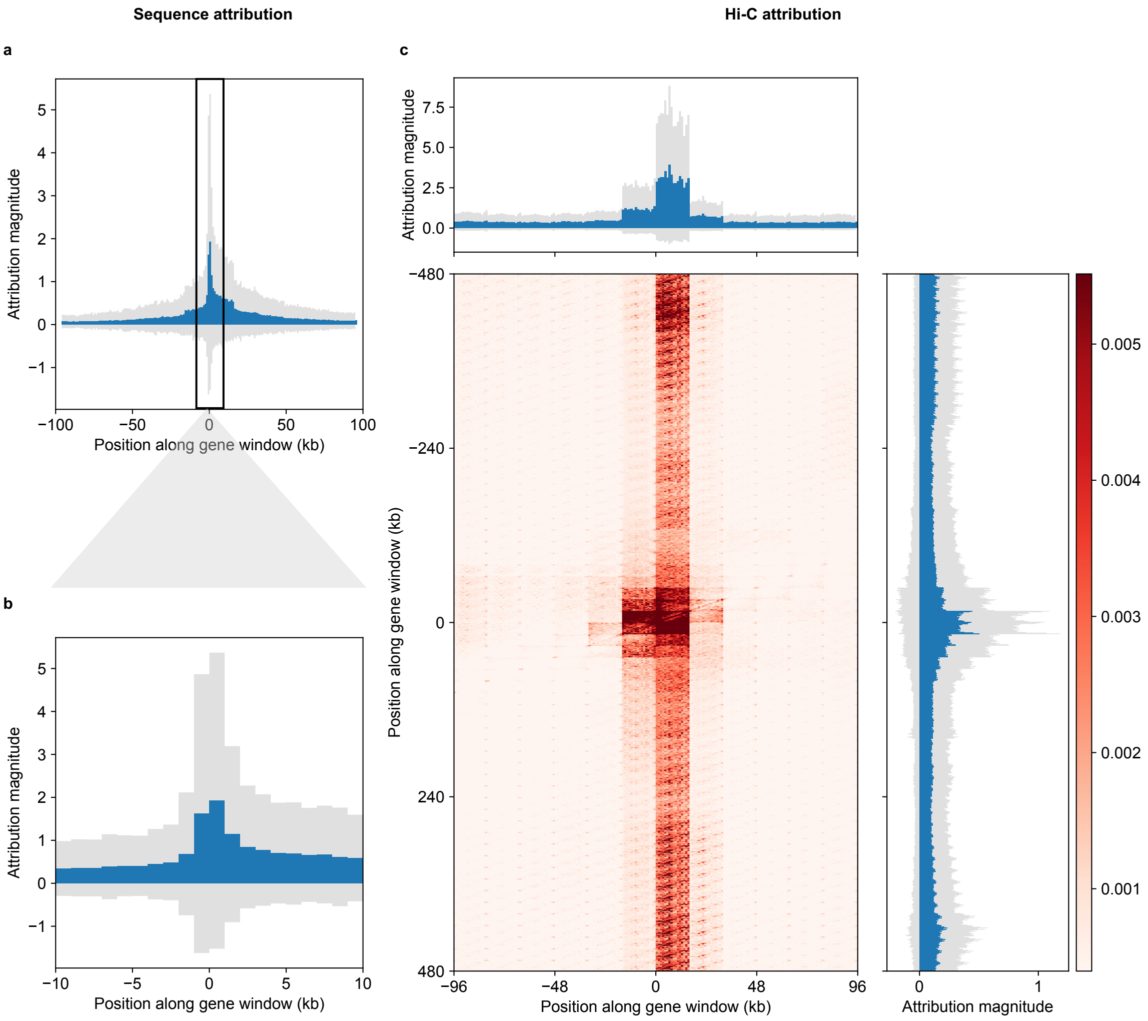


Figure S3: **Average sequence and Hi-C attribution maps for Puget.** Attributions are averaged across all test genes and all test biosamples (see Methods for attribution computation). Position 0 denotes the coordinate of the binned transcription start site (TSS), obtained by rounding the TSS coordinate down to the nearest 1,000 bp. **a** Average magnitude of sequence attribution across the 192 kb input window, aggregated into 1 kb bins. Blue bars show the mean attribution magnitude in each bin, and gray bands indicate one standard deviation. **b** Zoomed-in view of panel (a) around the binned TSS. Puget places the largest sequence attribution magnitude between $-1$ kb and $+1$ kb, consistent with a strong contribution from promoter-proximal sequence. **C** Average magnitude of Hi-C attribution represented as a TSS-centered 2D map at 1 kb $\times$ 1 kb resolution. The bottom left panel shows 2D attribution map. The top and right panels show 1D marginals obtained by summing the 2D map along the orthogonal axis; blue bars show the mean attribution magnitude and gray bands indicate one standard deviation. Hi-C attribution is most concentrated within $\pm16$ kb of the TSS along both axes, highlighting chromatin contacts involving promoters.

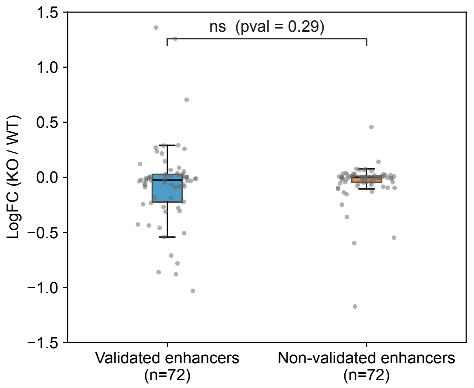


Figure S4: **Puget *in silico* perturbation effects for validated vs non-validated enhancer–gene pairs.** Distribution of Puget-predicted raw fold-change in gene expression after *in silico* perturbation of candidate enhancers, comparing validated versus non-validated enhancer–gene pairs from the K562 CRISPRi dataset. Because the dataset contains many more non-validated pairs, we construct a distance-matched negative set by subsampling non-validated pairs to match the distance-to-TSS distribution of validated pairs. Validated enhancers show more negative average effects than distance-matched non-validated enhancers, but the difference is non-significant.

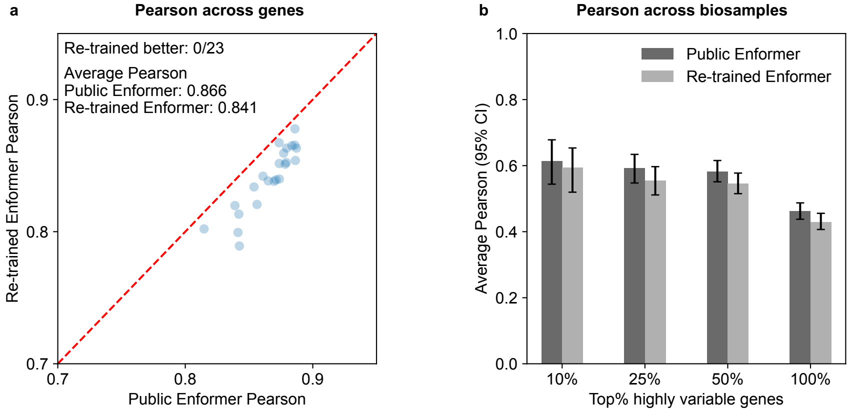


Figure S5: **Comparison between public Enformer and re-trained Enformer. a** Pairwise cross-gene performance comparing the public Enformer checkpoint with our re-trained Enformer. Across all 23 shared test chromosomes in the shared training biosamples, the re-trained Enformer underperforms the public Enformer, with the mean cross-gene Pearson correlation decreasing from 0.866 to 0.841. **b** Mean cross-biosample Pearson correlation as a function of the highly variable gene (HVG) threshold; across all thresholds, the re-trained Enformer performs comparably to the public Enformer. Error bars indicate 95% confidence intervals estimated from 1,000 bootstrap resamples of genes at each threshold.
