## supplementary-table-1 for "Puget predicts gene expression across cell types using sequence and 3D chromatin organization data"

Organism Biosource Experiment Type #Reads(M) #biological replicate split Date Lab Dataset Accession Hi-C Accession Organ Hi-C Donor RNA Donor RNA Date RNA Lab RNA Experiment Accession RNA Accession
human psoas muscle intact Hi-C 1405.10871 1 val 2022 "Erez Aiden, Baylor" ENCSR173QPK ENCFF625VNK muscle female adult (41 years) female adult (41 years) 2020 "Barbara Wold, Caltech" ENCSR574PFY ENCFF629LFN
human posterior vena cava intact Hi-C 952.050086 1 val 2022 "Erez Aiden, Baylor" ENCSR941AMF ENCFF946RZW heart female adult (59 years) female adult (59 years) 2020 "Barbara Wold, Caltech" ENCSR149AHS ENCFF658HZH
human left colon intact Hi-C 1771.34717 1 val 2022 "Erez Aiden, Baylor" ENCSR298UUL ENCFF035BLF colon female adult (59 years) female adult (59 years) 2020 "Barbara Wold, Caltech" ENCSR759TPN ENCFF635IYG
human PC-9 intact Hi-C 863.237325 1 val 2022 "Erez Aiden, Baylor" ENCSR859DRK ENCFF181ROW prostate   2021 "Barbara Wold, Caltech" ENCSR743GKS ENCFF877GJA
human PC-3 intact Hi-C 661.572043 2 val 2022 "Erez Aiden, Baylor" ENCSR038TZA ENCFF322IBI prostate   2020 "Barbara Wold, Caltech" ENCSR648KDM ENCFF617JTM
human OCI-LY7 intact Hi-C 1237.43839 1 val 2022 "Erez Aiden, Baylor" ENCSR088BOG ENCFF480KLP -   2021 "Barbara Wold, Caltech" ENCSR073XFZ ENCFF765ULK
human Caco-2 intact Hi-C 992.033677 1 val 2022 "Erez Aiden, Baylor" ENCSR820DYU ENCFF234MDO colon   2021 "Barbara Wold, Caltech" ENCSR469WPG ENCFF351WNJ
human spleen intact Hi-C 781.468918 1 train 2023 "Erez Aiden, Baylor" ENCSR692QWS ENCFF280EYD spleen female adult (61 years) female adult (61 years) 2020 "Barbara Wold, Caltech" ENCSR096LTX ENCFF143INW
human sciatic nerve intact Hi-C 805.197821 1 train 2022 "Erez Aiden, Baylor" ENCSR119TFS ENCFF602CHT nervous system female adult (41 years) female adult (41 years) 2020 "Barbara Wold, Caltech" ENCSR631NUQ ENCFF260VRG
human right lobe of liver intact Hi-C 1128.20982 1 train 2022 "Erez Aiden, Baylor" ENCSR753CKM ENCFF952JZV liver female adult (47 years) female adult (47 years) 2020 "Barbara Wold, Caltech" ENCSR323GUF ENCFF701OLG
human placenta intact Hi-C 811.814246 1 train 2022 "Erez Aiden, Baylor" ENCSR395DZW ENCFF490GGL placenta female embryo female embryo 2020 "Barbara Wold, Caltech" ENCSR899OKE ENCFF062KDX
human pancreas intact Hi-C 1855.12636 1 train 2022 "Erez Aiden, Baylor" ENCSR502GIO ENCFF586MQY pancreas female child (16 years) female child (16 years) 2020 "Barbara Wold, Caltech" ENCSR071DYD ENCFF258PUQ
human ovary intact Hi-C 1728.94203 1 train 2022 "Erez Aiden, Baylor" ENCSR782YIX ENCFF700CYI ovary female adult (59 years) female adult (59 years) 2020 "Barbara Wold, Caltech" ENCSR146ZLV ENCFF607DVO
human natural killer cell intact Hi-C 1404.07285 1 train 2022 "Erez Aiden, Baylor" ENCSR971CJS ENCFF197OWW blood male adult (33 years) male adult (33 years) 2022 "Barbara Wold, Caltech" ENCSR927KSI ENCFF909IYD
human mammary epithelial cell intact Hi-C 953.453127 1 train 2022 "Erez Aiden, Baylor" ENCSR707XVJ ENCFF512PQA breast female adult (19 years) female adult (19 years) 2021 "Barbara Wold, Caltech" ENCSR020YQE ENCFF292FVY
human lower lobe of left lung intact Hi-C 1413.32714 1 train 2022 "Erez Aiden, Baylor" ENCSR446NJD ENCFF395INO lung female adult (59 years) female adult (59 years) 2020 "Barbara Wold, Caltech" ENCSR718RTN ENCFF228JXX
human kidney intact Hi-C 1179.71168 1 train 2022 "Erez Aiden, Baylor" ENCSR044YKV ENCFF417GBZ kidney female adult (47 years) female adult (47

years) 2020 "Barbara Wold, Caltech" ENCSR892LBU ENCFF653OLW
human heart left ventricle intact Hi-C 2084.66314 1 train 2022 "Erez Aiden, Baylor" ENCSR324EJR ENCFF004YZQ heart male adult (55 years) male adult (54 years) 2023 "Barbara Wold, Caltech" ENCSR882RCG ENCFF789UOH
human endothelial cell of umbilical vein intact Hi-C 1106.35093 1 train 2022 "Erez Aiden, Baylor" ENCSR788FBI ENCFF783KQI uterus male newborn male newborn 2021 "Barbara Wold, Caltech" ENCSR993JMV ENCFF454MTF
human dorsolateral prefrontal cortex intact Hi-C 1445.70523 1 train 2022 "Erez Aiden, Baylor" ENCSR165UJN ENCFF925QIF brain male adult (78 years) male adult (78 years) 2022 "Barbara Wold, Caltech" ENCSR447WLU ENCFF354MPI
human colonic mucosa intact Hi-C 1338.75227 1 train 2022 "Erez Aiden, Baylor" ENCSR439ILZ ENCFF355NFJ colon female adult (41 years) female adult (41 years) 2020 "Barbara Wold, Caltech" ENCSR202OWR ENCFF169LQZ
human aorta intact Hi-C 1563.39569 1 train 2022 "Erez Aiden, Baylor" ENCSR554SLA ENCFF493YNC blood vessel female adult (59 years) female adult (59 years) 2020 "Barbara Wold, Caltech" ENCSR377FPC ENCFF568EVC
human activated T-helper 2 cell intact Hi-C 1476.96096 1 train 2022 "Erez Aiden, Baylor" ENCSR474VTW ENCFF687ATD spleen male adult (35 years) male adult (35 years) 2022 "Barbara Wold, Caltech" ENCSR341VFG ENCFF035GTI
human T-helper 2 cell intact Hi-C 1461.54203 1 train 2022 "Erez Aiden, Baylor" ENCSR832CKD ENCFF029UAL bone marrow male adult (35 years) male adult (35 years) 2022 "Barbara Wold, Caltech" ENCSR588TIV ENCFF279EMA
human "naive thymus-derived CD4-positive, alpha-beta T cell" intact Hi-C 1744.27501 1 train 2022 "Erez Aiden, Baylor" ENCSR215PTV ENCFF520GFL bone marrow male adult (35 years) male adult (35 years) 2022 "Barbara Wold, Caltech" ENCSR317HKT ENCFF138SCI
human MCF-7 intact Hi-C 846.34316 1 train 2022 "Erez Aiden, Baylor" ENCSR660LPJ ENCFF420JTA breast 2020 "Barbara Wold, Caltech" ENCSR355JZC ENCFF721BRA
human MCF 10A intact Hi-C 699.302541 2 train 2022 "Erez Aiden, Baylor" ENCSR370TFL ENCFF977XWK breast 2021 "Barbara Wold, Caltech" ENCSR815NTL ENCFF215WRT
human HCT116 intact Hi-C 1466.22 2 train 2022 "Erez Aiden, Baylor" ENCSR477GZK ENCFF573OPJ colon 2020 "Barbara Wold, Caltech" ENCSR698RPL ENCFF435PHM
human Calu3 intact Hi-C 900.045158 1 train 2022 "Erez Aiden, Baylor" ENCSR118AQZ ENCFF127TPS lung 2021 "Barbara Wold, Caltech" ENCSR897JEH ENCFF472SUX
human A673 intact Hi-C 1393.27168 1 train 2022 "Erez Aiden, Baylor" ENCSR455KLX ENCFF129LMU bone 2021 "Barbara Wold, Caltech" ENCSR558SEE ENCFF619BUB
human Panc1 intact Hi-C 901.48905 1 test 2022 "Erez Aiden, Baylor" ENCSR584RBV ENCFF522YLZ pancreas 2020 "Barbara Wold, Caltech" ENCSR128CYL ENCFF710IFD
human K562 intact Hi-C 999.493934 2 test 2022 "Erez Aiden, Baylor" ENCSR479XDG ENCFF621AIY bone marrow 2020 "Barbara Wold, Caltech" ENCSR792OIJ ENCFF928NYA
human IMR-90 intact Hi-C 1085.83475 2 test 2022 "Erez Aiden, Baylor" ENCSR345VTI ENCFF281ILS lung 2020 "Barbara Wold, Caltech" ENCSR797RXV ENCFF325KTI
human HepG2 intact Hi-C 653.713349 1 test 2022 "Erez Aiden, Baylor" ENCSR888DEJ ENCFF807IRK liver 2020 "Barbara Wold, Caltech" ENCSR245ATJ ENCFF863QWG
human GM12878 intact Hi-C 1756.29724 1 test 2022 "Erez Aiden, Baylor" ENCSR916MFV ENCFF318GOM blood 2020 "Barbara Wold, Caltech" ENCSR820PHH ENCFF345SHY

human B cell intact Hi-C 1358.44992 1 test 2022 "Erez Aiden, Baylor" ENCSR847RHU ENCFF076LWH bone marrow male adult (22 years) male adult (22 years) 2022 "Barbara Wold, Caltech" ENCSR896YYL ENCFF192WQE
mouse adrenal gland intact Hi-C 1519.67 1 test 2025 "Erez Aiden, Baylor" ENCSR820CZQ ENCFF417VZK adrenal gland B6XCast EiJ_Female_2m B6CASTF1/J female adult (2 months) 2021 "Barbara Wold, Caltech" ENCSR288BJQ ENCFF883HID
mouse gastrocnemius intact Hi-C 1580.04 1 test 2025 "Erez Aiden, Baylor" ENCSR622SLO ENCFF371FGB muscle B6XCast EiJ_Female_2m B6CASTF1/J female adult (2 months) 2021 "Barbara Wold, Caltech" ENCSR791UVS ENCFF731ZHP
mouse left cerebral cortex intact Hi-C 2139.99 1 test 2025 "Erez Aiden, Baylor" ENCSR639NRL ENCFF969SFG left cerebral cortex B6XCast EiJ_Female_2m B6CASTF1/J female adult (2 months) 2021 "Barbara Wold, Caltech" ENCSR219ZXZ ENCFF806DOB
mouse heart intact Hi-C 1864.19 1 test 2025 "Erez Aiden, Baylor" ENCSR997TTD ENCFF651SGY heart B6XCast EiJ_Female_2m B6CASTF1/J female adult (2 months) 2021 "Barbara Wold, Caltech" ENCSR984JDC ENCFF894UGW
